# Reaching around obstacles accounts for uncertainty in coordinate transformations

**DOI:** 10.1101/706317

**Authors:** Parisa Abedi Khoozani, Dimitris Voudouris, Gunnar Blohm, Katja Fiehler

**Affiliations:** Centre for Neuroscience Studies, Queen’s University, Kingston, Ontario, Canada; Canadian Action and Perception Network (CAPnet), Toronto, Ontario, Canada; Association for Canadian Neuroinformatics and Computational Neuroscience (CNCN), Kingston, Ontario, Canada; Center for Mind, Brain, and Behaviour, Marburg University, Marburg, Germany; Psychology and Sport Sciences, Justus Liebig University, Giessen, Germany

## Abstract

When reaching to a visual target, humans need to transform the spatial target representation into the coordinate system of their moving arm. It has been shown that increased uncertainty in such coordinate transformations, for instance when the head is rolled toward one shoulder, leads to higher movement variability and influence movement decisions. However, it is unknown whether the brain incorporates such added variability in planning and executing movements. We designed an obstacle avoidance task in which participants had to reach with or without visual feedback of the hand to a visual target while avoiding collisions with an obstacle. We varied coordinate transformation uncertainty by varying head roll (straight, 30° clockwise and 30° counterclockwise). In agreement with previous studies, we observed that the reaching variability increased when the head was tilted. Indeed, head roll did not influence the number of collisions during reaching compared to the head straight condition, but it did systematically change the obstacle avoidance behavior. Participants changed the preferred direction of passing the obstacle and increased the safety margins indicated by stronger movement curvature. These results suggest that the brain takes the added movement variability during head roll into account and compensates for it by adjusting the reaching trajectories.

## Introduction

Transforming retinal information to the coordinate system of the moving arm is crucial for performing visually guided movements, e.g. reaching (Buneo & Andersen, 2006; Buneo et al., 2002; Cohen & Andersen, 2002; Engel, Flanders, & Soechting, 2002; Knudsen, 2002; Knudsen, du Lac, & Esterly, 1987; Lacquaniti & Caminiti, 2003; Soechting & Flanders, 1992). It has been suggested that coordinate transformations should be considered as stochastic processes (e.g. as processes causing random signal-dependent noise in the transformed signal) that add uncertainty to the transformed signals (Alikhanian, Carvalho, & Blohm, 2015; McGuire & Sabes, 2009; Sober & Sabes, 2003, 2005). Furthermore, it has been shown that stochasticity in coordinate transformations propagates to the movement resulting in increased movement variability (Abedi Khoozani & Blohm, 2018; Burns, Nashed, & Blohm, 2011; Burns & Blohm, 2010; Schlicht & Schrater, 2007); however, it is unknown if the brain accounts for potential consequences of such added movement variability while planning and executing reaching movements.

Accurate coordinate transformations rely on the estimation of three-dimensional (3D) body pose (Blohm & Crawford 2007). This requires an internal model of different body parts with regard to each other, e.g. eye relative to head translation, and an estimation of joint angles, e.g. head rotation. While internal models are learned and resistant to change, the estimation of the joint angle can arise from two sources: 1) afferent sensory signals and 2) efferent copies of motor commands. Both sources are corrupted with uncertainty in sensory processing and variability of neuronal spiking (Faisal, Selen, & Wolpert, 2009). Several studies have suggested that varying body pose, e.g. rolling the head, increases movement variability (Abedi Khoozani & Blohm, 2018; Burns, Nashed, & Blohm, 2011; Burns & Blohm, 2010; Schlicht & Schrater, 2007). For instance, Burns and Blohm (2010) showed that rolling the head to either shoulder results in higher goal-directed reaching variability compared to straight head reaching. The authors argued that this increased variability stems from the signal-dependent noise during the required sensory estimations of body geometry (here, head angle) that are necessary for accurate coordinate transformations. However, another interpretation can be that since humans perform reaching mostly with the head in an upright posture, the difference in variability can arise from the lack of experience, or less familiarity, in the rolled condition (Sober & Körding, 2012).

To differentiate between these two speculations, we have previously asked humans to perform visually guided reaching movements while their heads were rolled or their necks loaded with an external mass (Abedi Khoozani & Blohm, 2018). Our rationale was that if lack of familiarity caused the added variability, then neck load should have no effect; conversely, active estimation of head angles should result in larger variability. This higher variability stems from the signal-dependent noise due to increased muscle activity (Cordo et al., 2002; Faisal et al., 2008; Lechner-Steinleitner, 1978; Sadeghi et al., 2007; Scott & Loeb, 1994) induced by neck load. Since larger joint angle estimates and muscle activations are accompanied with higher uncertainty (Blohm & Crawford, 2007; Van Beuzekom & Van Gisbergen, 2000; Wade & Curthoys, 1997), both head roll and neck load manipulations should result in noisier coordinate transformations. Our result supported the hypothesis that signal-dependent noise in coordinate transformations increases movement variability (Abedi Khoozani & Blohm, 2018). Additionally, we observed that both rolling the head and loading the neck results in angular reaching biases. Using our computational model, we showed that these biases can be explained by over- and under-estimation of sensed head angles compared to actual head angles. Based on these studies, we concluded that biases and uncertainties associated with head angle estimation during reaching propagate to the coordinate transformations resulting in added biases and variability in reaching movements. However, it is unknown if the brain incorporates this added movement variability when planning and executing reaching movements.

One approach to investigate whether the brain is accounting for the added movement variability caused by stochastic coordinate transformations is to perform reaching movements in constrained environments, i.e. in the presence of obstacles. A failure in accounting for the added variability should result in behavioral consequences, i.e. obstacle collisions. In general, humans are successful in avoiding obstacles and they do so by accounting for several factors such as sensory uncertainty (Cohen, Biddle, & Rosenbaum, 2010), motor noise (Cohen et al., 2010; Hamilton & Wolpert, 2002), and biomechanical costs (Cohen et al., 2010; Sabes, 1997; Sabes, Jordan, & Wolpert, 1998; Voudouris, Smeets, & Brenner, 2012). For instance, Cohen et al. (2010) showed that both higher visual uncertainty and increased motor noise resulted in increased distance from the obstacle (increased safety margins). Therefore, based on these studies humans are capable to account and compensate for the increased motor noise. Thus, we chose an obstacle avoidance task as it provides a suitable test bed to evaluate if increased noise induced by stochastic coordinate transformations is considered during reaching.

To investigate whether humans can compensate for the higher movement variability caused by stochastic coordinate transformations, we designed a reaching task to a visual target while avoiding an obstacle. For modulating the uncertainty in coordinate transformations, participants performed the reaching movements with different head rolls (30° toward right shoulder (clockwise; CW), 0°, and 30° toward the left shoulder (counterclockwise; CCW)). Following previous studies, we expected that varying head roll results in higher movement variability and rotational biases (Abedi Khoozani & Blohm, 2018; Alikhanian et al., 2015; Burns & Blohm, 2010). If the brain does not consider the added movement variability for reaching, we hypothesize higher collision rates in the rolled head compared to the straight head condition. On the other hand, if the brain considers the added movement variability, the collision rates should be similar for all head roll conditions, which would consequently come with adapted movement strategies to compensate for the added movement variability (e.g. decreased movement speed, changed preferred direction of passing the obstacle, and/or increased safety margins). Furthermore, previous studies showed that providing visual feedback of the moving hand will alleviate the effect of stochastic coordinate transformations (Blohm & Crawford, 2007). Therefore, we asked participants to perform reaching movements with or without visual feedback. We expect lower compensation, if there is any, for the visual feedback condition compared to the no-visual feedback condition. In agreement with previous studies (Abedi Khoozani & Blohm, 2018; Alikhanian et al., 2015; Burns & Blohm, 2010), we observed that movement variability increased when the head was tilted. In both feedback conditions, this was accompanied by a change in the preferred direction of passing the obstacle and increased safety margins, while the collision rate remained unaffected. We conclude that the brain accounts for the added uncertainty due to coordinate transformations and compensates for it by increasing safety margins whenever task performance is compromised.

## Materials and methods

### Participants

We collected data from 18 healthy humans (10 female) aged between 19 to 38 years (M = 25 years) with normal or corrected to normal vision. All participants were right handed by self-report, and free of any known neurological issues. The experiment was approved by the Justus Liebig University Giessen, general board of ethics and all participants gave their written consents. They received monetary compensation (8 € / hour) or course credits for their participation.

### Apparatus and Task

Participants were seated in front of a workspace that comprised a robotic setup with a graspable handle (vBot; Howard, Ingram, & Wolpert, 2009), a monitor, and a mirror. A helmet with a protruding long stick and a measuring framework were used to control for the head roll in each condition (Figure 1A). Visual stimuli were presented on the monitor and reflected in the mirror, which was placed above the robot (Figure 1B).

**Figure 1.**
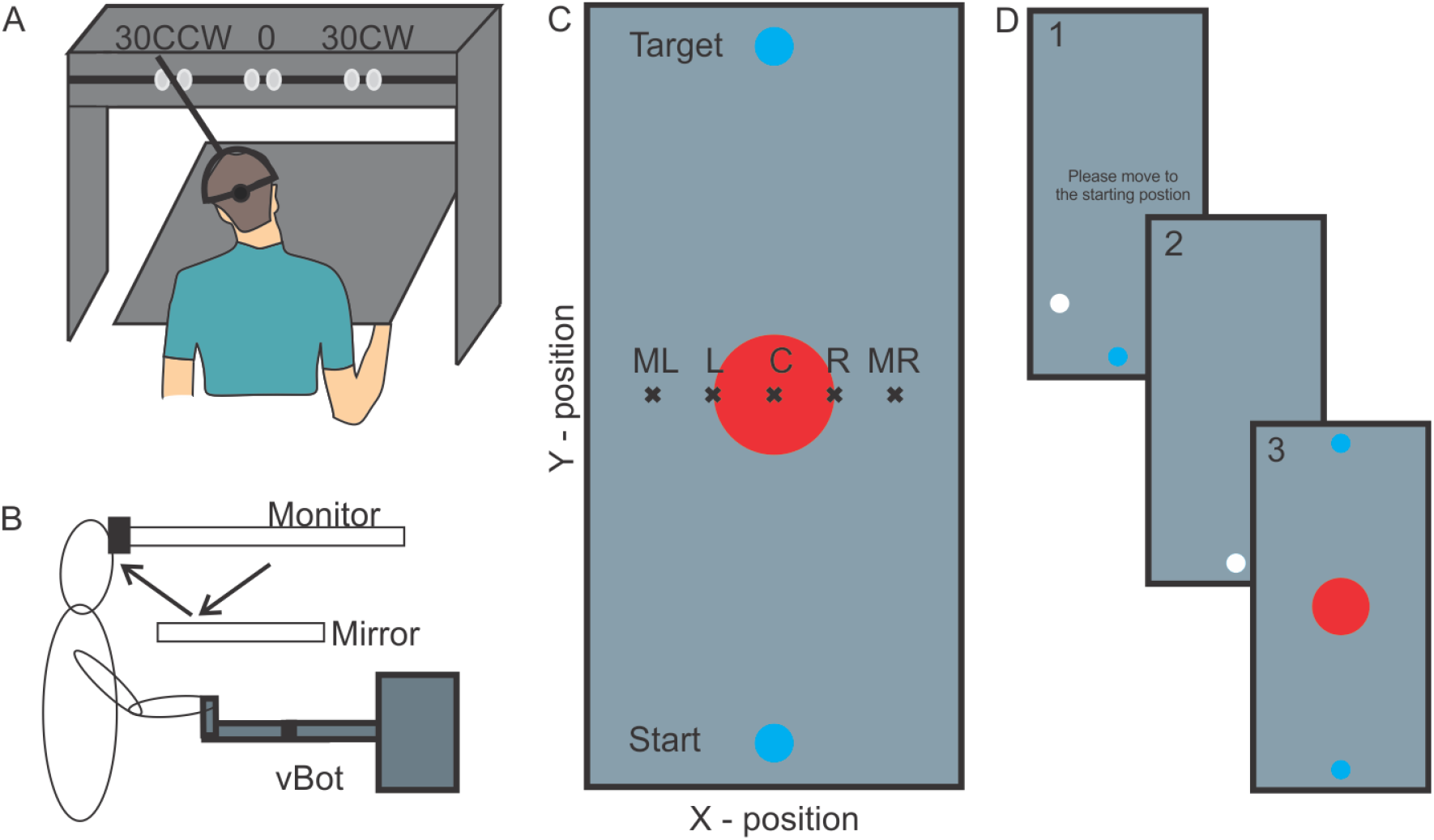
Overview of the experimental setup. A) Participants performed the reaching task in three head orientations (30° CCW, 0, and 30° CW). Here only the framework of the robotic setup and the head orientation setup is displayed, B) The vBot setup consisted of a mirror that displayed the task instructions from the monitor. Below the mirror, a robot handle was placed that could be freely moved by the participants. C) Participants brought their hand to the start point and reached to the target while avoiding hitting the obstacle in the center. Obstacles were randomly shifted to the left or right, along the X-position, across trials. Possible shift positions are depicted by “×” with the letters indicating the shifts; ML: most leftward, L: leftward, C: central, R: rightward, MR: most rightward, D) Example trial: first, participants were instructed to bring their hand to the starting position (1), as soon as they arrived at the starting position (2), the target position and the obstacle (here central) appeared and participants were instructed to move to the target position in less than 1000 ms (3).

A mirror was placed horizontally in front of the participants. This mirror prevented vision of the arm so that participants were unable to see the movements that they performed below the mirror. The visual stimuli were presented on a monitor and reflected on the mirror that was placed below it (Figure 1B). Four disks and the visual instructions were presented in each trial. Two discs (0.5 cm diameter) were blue, each serving as starting and target position. Both were located in the middle of the screen laterally (X-position; which was the same as the body midline). The starting and target positions were 9 and 11 cm closer and further from the centre of the screen, respectively, resulting in a distance of 20 cm between the two. A third disc (1.8 cm diameter and red color) served as the obstacle. It was presented in the middle of the screen and in five possible positions: at the center (in line with the starting and target position) or shifted by 1.8 cm or 3.6 cm either to the right or left of that central position (Figure 1C). To simulate a physical obstacle, a repulsive force-field (between 0 to 40N, faster movements toward the center resulted in higher force) from the center of the obstacle was applied (i.e. vBotDisc). Consequently, the handle could not move into the obstacle. Finally, a fourth, white disk (0.5 cm diameter) represented the position of the robotic handle, whenever visual feedback was provided. A visual instruction that was prompting participants to move to the starting position at the beginning of the trial was presented at the centre of the screen. Visual stimuli were implemented using Psychtoolbox (MATLAB 2015) while C++ programming was used for implementing the force-field and programming the robot.

In order to reach to the viewed target, participants grasped the robot’s handle with their right hand while resting their forehead on the workspace’s framework in front of the screen. They performed the task in three possible head orientations: 30° CCW, 0°, 30° CW. Each head orientation condition was performed with and without visual feedback of their reaching hand. The visual feedback was provided by a moving white disc (0.5 cm diameter) on the screen representing the robot handle position. This resulted in six combinations, each of which was presented in separate blocks of trials. At the beginning of each block, the experimenter positioned the participant’s head in the respective orientation. Before the start of each trial, participants grasped the robot handle and brought it to a fixed starting position using visual feedback of the robot’s handle. After positioning the hand on the starting position, the target and an obstacle were simultaneously displayed on the mirror. Participants were instructed to immediately start reaching towards the visual target while avoiding the obstacle. The trial was ended as soon as the participants’ hand distance from the center of the target was less than 0.5 cm when hand visual feedback was provided. When visual feedback of the hand was absent, the end of the trial was defined as the moment when the participants’ hand reached the position of the target in depth of the target and was less than 4 cm away from its lateral position. If participants were able to reach the target without hitting the obstacle in less than 1000 ms from the moment of target presentation, the visual target would turn green indicating that the trial was successful. Otherwise the target would turn red indicating that the trial was aborted and would be repeated later. At the end of each trial all visual stimuli disappeared and the next trial started with the appearance of the starting position.

Before starting the experiment, participants performed a short practice block with 20 trials. Within each of the six blocks, each obstacle position was presented 48 times, resulting in a total of 240 trials per block, and thus in a total of 1440 trials for the complete experiment, which lasted roughly 60 minutes. The combination of the head angle and visual feedback for each block was chosen based on Latin squares method to counterbalance among all conditions (Jacobson & Matthews, 1998).

### Data analysis

All offline analyses were performed using MATLAB 2018. To test whether reaching movement strategies are altered for different head roll conditions, we required to create a reliable and normalized estimate of the trajectory for each trial using functional data analysis (Ramsay & Silverman, 2005), which fits each dimension of the raw data (x and y) with B-splines. These spline functions are good candidates to fit motion data that are not strictly periodic (to see studies using similar techniques see Gallivan & Chapman, 2014; Loehr and Parlmer, 2007). We fitted order 6 splines to each dimension of the data. Since our trajectory data did not include missing points, we chose to have 10 equally distributed breakpoints across the data. To perform the data fits, we used ‘splinefit.m’ in MATLAB 2018a. Using this technique, we were able to create a continuous representation of our data for each dimension which is scale invariant. Therefore, we could extract as many points in time or space as desired without distorting the spatiotemporal features of the movement trajectories. To verify that the difference between re-sampled trajectories and recorded trajectory is not significant, we compared the two trajectories. To do so, we calculated the mean squared errors between the two trajectories for each participant and observed that the difference between the two trajectory is negligible (mean squared error = 2.5 ± 5.6 mm; for more details refer to Supplementary figure). For the analysis, we sampled 2000 equally-spaced time points from the fitted spline functions. As it is demonstrated in previous work (Whitwell and Goodale, 2013), the matter of normalization is a critical choice. Typically, normalization should be performed along the dimension which is not varying due to the experimental conditions. In our experiment, participants were constrained in performing the task under 1 second. Therefore, we chose time as the normalization dimension. To make sure that the normalization method did not affect our final result, we calculated the reaction time and movement time and assessed if the experimental conditions significantly affected these temporal parameters.

A central differential algorithm (differentiation was performed in a monotonically time increasing manner) was used to calculate hand velocity and acceleration. Before each derivation, a low-pass filter (autoregressive forward-backward filter, cutoff frequency = 50 Hz) was used to smooth the data. We calculated the movement onset by finding the moment of 25% and 75% of the trial’s peak velocity and then extrapolating a line between these two moments until this line crossed that trial’s baseline velocity as this was measured by averaging the velocity during the first 200 ms of the trial (for futher details see Brenner & Smeets, 2018). The reaction time (RT) was calculated by subtracting the time of movement onset from the time that the target appeared. The movement duration (MD) was calculated as the time difference between the end of the trial (see Apparatus and Task) and movement onset. Trials with RT < 100 ms (predictive movements) and RT+MD > 1000 ms (too slow) were removed from the analysis (1.7%) and considered as invalid trials.

To assess the effect of varying head orientation on movement behavior, we calculated movement variability across the whole reaching trajectory. To do so, the normalized trajectories of each participant were averaged separately for each obstacle position and movement direction (rightward or leftward of that obstacle). Since movements were predominantly along the Y-position direction, we only calculated the standard deviation of the handle’s lateral position across trials for each of the 2000 normalized steps. Then, we calculated the boundaries of the averaged trajectories by adding/subtracting the standard deviation to/from the mean of the trajectory along the X-position (Figure 2A). Finally, we calculated the movement variability as the area between the trajectory boundaries. We performed these steps separately for each participant, head orientation, visual condition, and movement direction for each obstacle position. We considered the calculated movement variability for different directions of each obstacle as valid only if the number of movements in a certain direction for a given obstacle was more than 10% of the overall movements for that obstacle.

**Figure 2.**
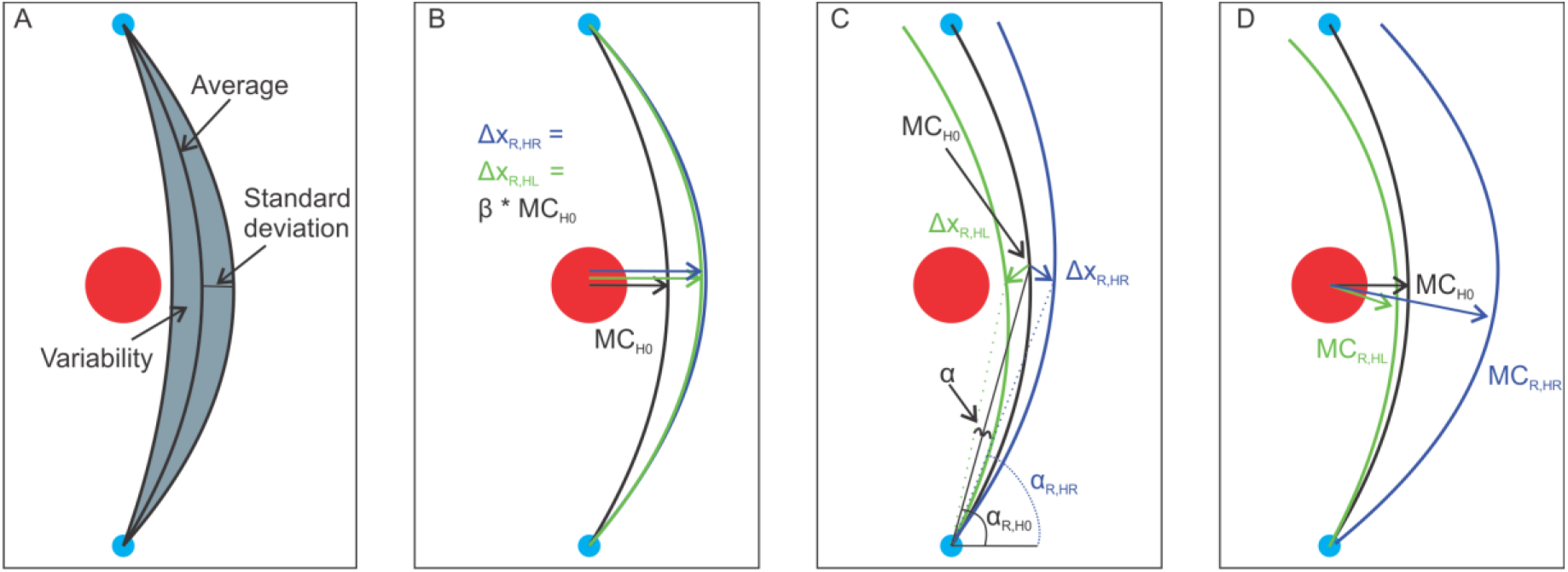
Variability, expansion, and rotational biases calculations. A) The overall movement variability was calculated as the area between the movement boundaries. The boundaries were calculated for each participant separately by adding/subtracting the standard deviation of the lateral movements from the average for all 2000 samples along the trajectory, B) expansion biases (β): varying head orientation increases movement variability resulting in hand trajectories moving away from the obstacle, C) rotational biases (α): we hypothesized that rolling the head creates rotational biases resulting in symmetrical shifts of the trajectories for the 30° CW/CCW head orientations (colored solid lines) around the straight head (black solid line), D) combination of rotational biases and expansion biases result in an asymmetry in the shifted trajectories for the 30° CW/CCW head orientations (colored solid lines) compared to the straight head (black solid line).

To investigate if participants were able to compensate for the effect of varying head roll, we calculated the collision rate via dividing the number of collisions with the obstacle by the total number of valid trials for each head roll and visual feedback condition. In the next step, we assessed whether the added movement variability due to rolling the head had a tangible effect on the movement strategies. To do so, we considered two parameters: the preferred direction of passing the obstacle (i.e. around the right or the left side of the obstacle) and the safety margins (i.e. distance from the obstacle at the moment of passing it). For direction of passing the obstacle, we calculated the percentage of rightward movements and expected to see higher rightward movements for when the obstacle was shifted to the left and the reverse for when the obstacle was shifted to the right. We expected similar percentage of rightward and leftward movements for the central obstacle. We hypothesized that rolling the head should modulate the preferred direction of passing the obstacle to decrease the likelihood of collision; most noticeably for the central obstacle. With regards to safety margins, we hypothesized that reaching trajectories should deviate further away from the obstacle in the rolled conditions in order to compensate for the expected added movement variability. This should be reflected in a larger curvature which is more noticeable for the central obstacle (β: expansion biases; Figure 2B). However, based on our earlier findings and due to under- or over-compensation for head roll (Abedi Khoozani & Blohm, 2018), we further expected that movement trajectories for straight head conditions should fall symmetrically between the trajectories for the two head roll conditions (α: rotational biases; Figure 2C). In our data, both expansion and rotational biases are combined. Consequently, a simple analysis of the curvature for different head roll condition is not revealing all necessary information for the evaluation of the chosen movement strategies.

To separate rotational and expansion biases from each other we employed the following method. First, we considered the straight head condition as baseline (no effect of head roll is expected). Since we are interested in extracting the expansion biases, we picked the point with the maximum possible effect: maximum curvature. To quantify the overall effect of head roll we calculated the difference of the maximum curvature of the averaged trajectories between straight head and 30° CW/CCW head orientations (Δ*x*). As it is demonstrated in Figure 2 B-C, expansion and rotational biases affect the trajectory differently. That is, expansion biases should cause the shift in curvature in the same direction:

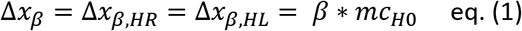

Where the first subscript indicates the biases and the second subscript indicates the head orientation (HL: 30°CCW and HR: 30°CW).

On the other hand, rotational biases are expected to be symmetrical around control condition:

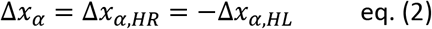

By the assumption that expansion and rotational biases are added together, we have:

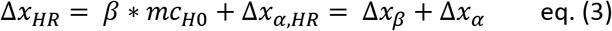

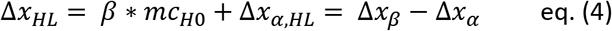

The visualization of the variables for the rightward movement direction is provided in Figure 2D.

Consequently, we can separate the shifts in trajectories caused by expansion (Δ*x*_*β*_) and rotation (Δ*x*_*α*_) using the following calculation:

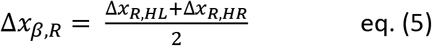

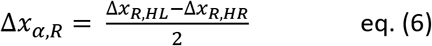

Where the first subscripts indicate the movement direction (R: rightward and L: leftward). Thus, we calculated the percentage of expansion as following:

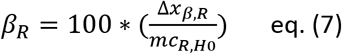

Therefore, positive values indicate expansion while negative values indicate shrinkage. Similar calculations can be applied for the leftward movements.

### Statistical analysis

We used JASP (https://jasp-stats.org/) to perform the statistical analyses. To examine the effect of head orientation (0 and 30° CW/CCW), obstacle position (most leftward, leftward, central, rightward, and most rightward), and visual feedback (with and without) on the above-mentioned dependent variables (movement variability, collision rate, movement speed, preferred direction of passing the obstacle, and safety margins (calculated as expansion biases)), we deployed repeated measures ANOVA or student t-test. The exact test and design was identified based on the question of interest. We provide this information in individual sections in the Results to avoid repetition. Significant differences between the conditions were further investigated with two-sample paired t-tests and the reported p-values were Bonferroni-Holm corrected.

### Functional comparisons of the trajectory data

Similar to the approach reported by Gallivan and Chapman (2014), we performed functional analysis of variance (fANOVA) on normalized trajectories. A fANOVA is an extension of the traditional ANOVA that can be applied on continuous data. More technically, fANOVA provides F-statistics along the normalized axes (here time) that shows where the trajectories deviate significantly from each other for different conditions. When only two trajectories are compared, fANOVA acts as a functional t-test. We first deployed fANOVA to assess if varying head rolls result in deviation of trajectories from each other. If we found the main effect of the head orientation on trajectories to be significant, then, we performed the functional t-test to assess where the trajectories for each head roll condition deviated from the straight head condition.

To perform the above-mentioned analysis, we deployed the custom MATLAB algorithms developed by Gallivan and Chapman (2014). We should note that based on the designed algorithm we needed to down sample our trajectories to 200 points. We performed this by resampling from the fitted Splines.

The data and analysis codes are provided online. The link can be found in the Endnote section.

## Results

The objective of this study was to investigate whether humans compensate for movement variability caused by stochastic coordinate transformations. To this aim, we asked participants to reach to a visual target without colliding with an obstacle while having different head rolls (30° CW/CCW and 0°). Figure 3 illustrates the trajectories of two example participants (#2 and #16). Both participants were able to successfully avoid the obstacle in all head orientations (the collision rate increase was less than 2%), however, each of the two participants showed a different movement behavior in the rolled head orientations (green and blue) compared to the straight head (black). Specifically, participant #2 moved further away from the obstacle to successfully reach to the target (increased safety margin). In contrary, participant #16 kept the same distance from the obstacle, but instead decreased the movement speed. Based on these results, humans seem to change their movement strategy for different head orientations.

**Figure 3.**
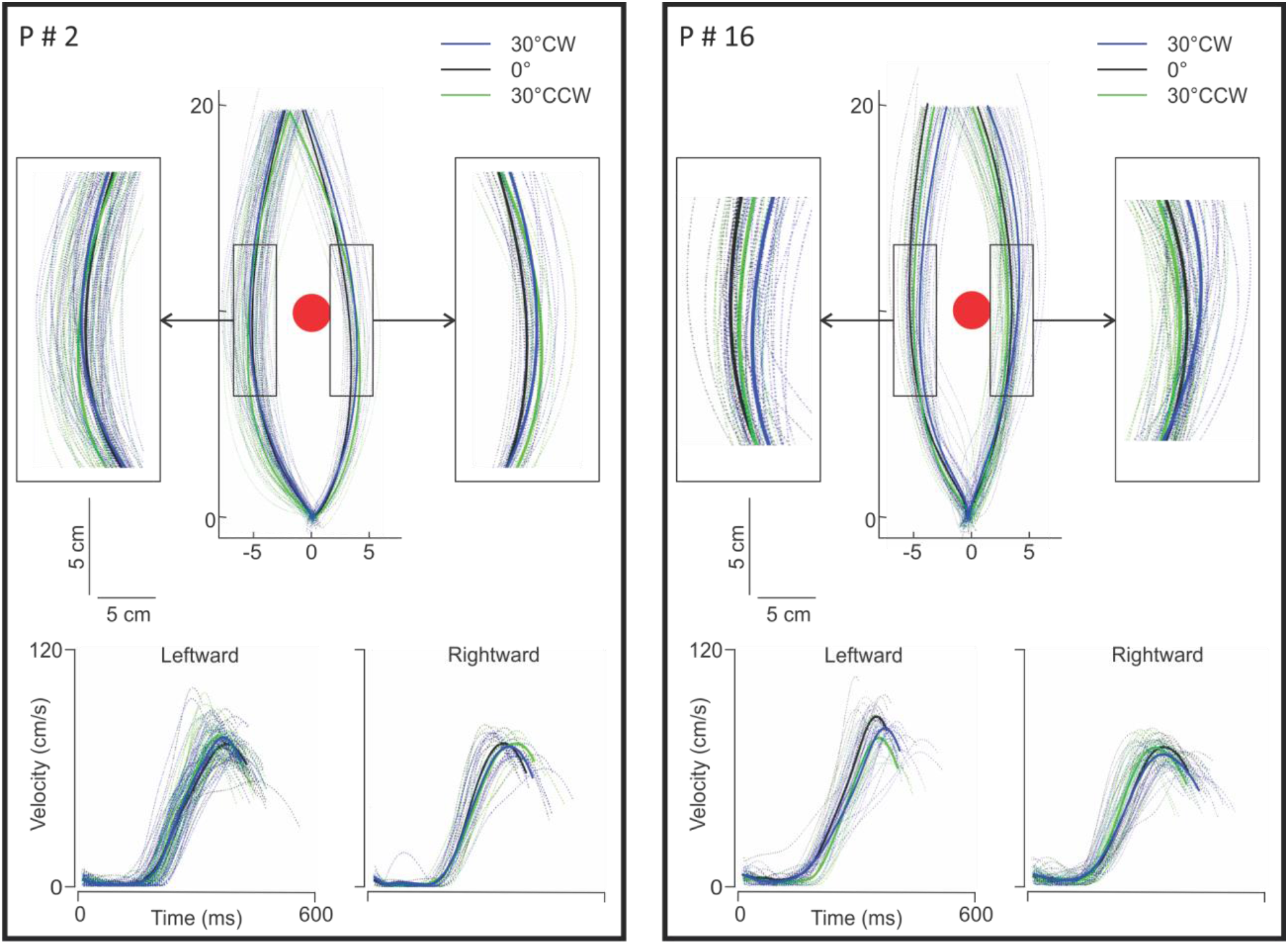
Obstacle avoidance strategies of two participants. The data is shown for the central obstacle position (red circle) without visual feedback. The left panel illustrates the trajectory data as well as the velocity data for participant #2. This participant moved further away from the obstacle, specifically for the rightward movements, in the head roll conditions (green and blue solid lines) compared to straight head condition (solid black line). The two panels on the sides illustrate a zoomed version of the trajectories. The peak velocity did not change for the rightward movements while increased for the leftward movements. The right panel illustrates the behavior of participant #16. In contrary to participant #2, this participant decreased the peak velocity and also decreased the distance from the obstacle, especially for leftward movements.

**Figure 4.**
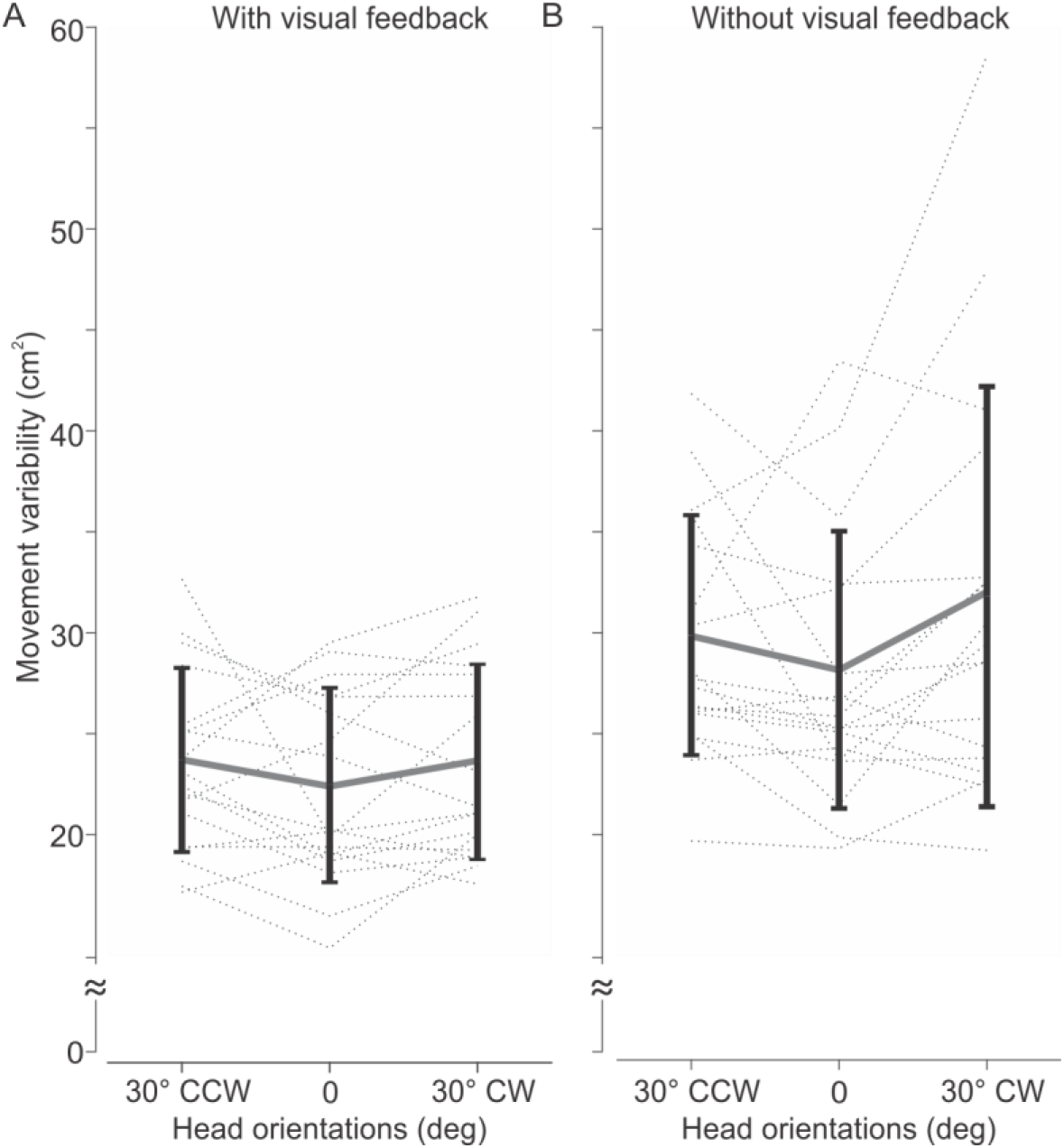
Effect of varying head orientation on movement variability. A) Visual feedback condition: varying the head orientation did not affect the movement variability. B) Without visual feedback condition: participants showed different effects of varying head orientation on their movement variability. Some participants showed increased movement variability while others showed decreased movement variability. Error bars are standard deviations.

### Rolling the head increased movement variability

Previous studies demonstrated that rolling the head while reaching increases movement variability (Burns & Blohm, 2010; Abedi Khoozani & Blohm, 2018). Therefore, in the first step we investigated the effect of varying head orientation on movement variability depending on the visual feedback of the hand. To remind the reader, we calculated the movement variability as the surrounding area between the lateral deviations from the averaged trajectory. Since we did not expect to observe any effect of obstacle position on the movement variability, we performed a 3 (head orientation) × 2 (visual feedback) repeated measures ANOVA. We observed a main effect of head roll (F(2,34)=4.39, p = 0.020, η^2^=0.205), a main effect of visual feedback (F(1,17)=31.36, p = 3.190e-5, η^2^=0.648) and no interaction between head roll and visual feedback (F(2,34)=14.08, p = 0.403). Overall, movement variability was larger when visual feedback of the hand was removed. Post-hoc t-tests for the head orientation effect revealed a significant increase of movement variability for the CW head orientation compared to straight head (t(17) = 2.729, p = 0.043, Cohen’s d = − 0.643), and a trend for the CCW head orientation compared to straight head (t(17) = 2.171, p = 0.089). Thus, these results confirm previous work that rolling the head increases the movement variability.

### Increased movement variability did not affect the collision rate for different head orientations

If the brain does not consider the added movement variability caused by stochastic coordinate transformations, collision rates should be higher for the rolled (CW, CCW) than the straight head conditions. As we did not expect to observe any difference in the collision rate for the different obstacle positions, we pooled the data across the obstacle positions and assessed if head orientation or visual feedback affected the collision rate. The 3 (head orientation) × 2 (visual feedback) repeated measures ANOVA revealed no main effect of head orientation (F(2,34) = 0.100, p = 0.905, η^2^=0.006), a main effect of visual feedback (F(1,17) = 12.831, p = 0.002, η^2^=0.430), and no interaction between the two (F(2,34) = 1.044, p = 0.363, η^2^=0.058). As illustrated in Figure 5, removing visual feedback caused an increase in the collision rate. However, in both visual conditions, the collision rate remained the same for different head orientations indicating that participants were able to successfully compensate for the added variability due to varying head orientations.

**Figure 5.**
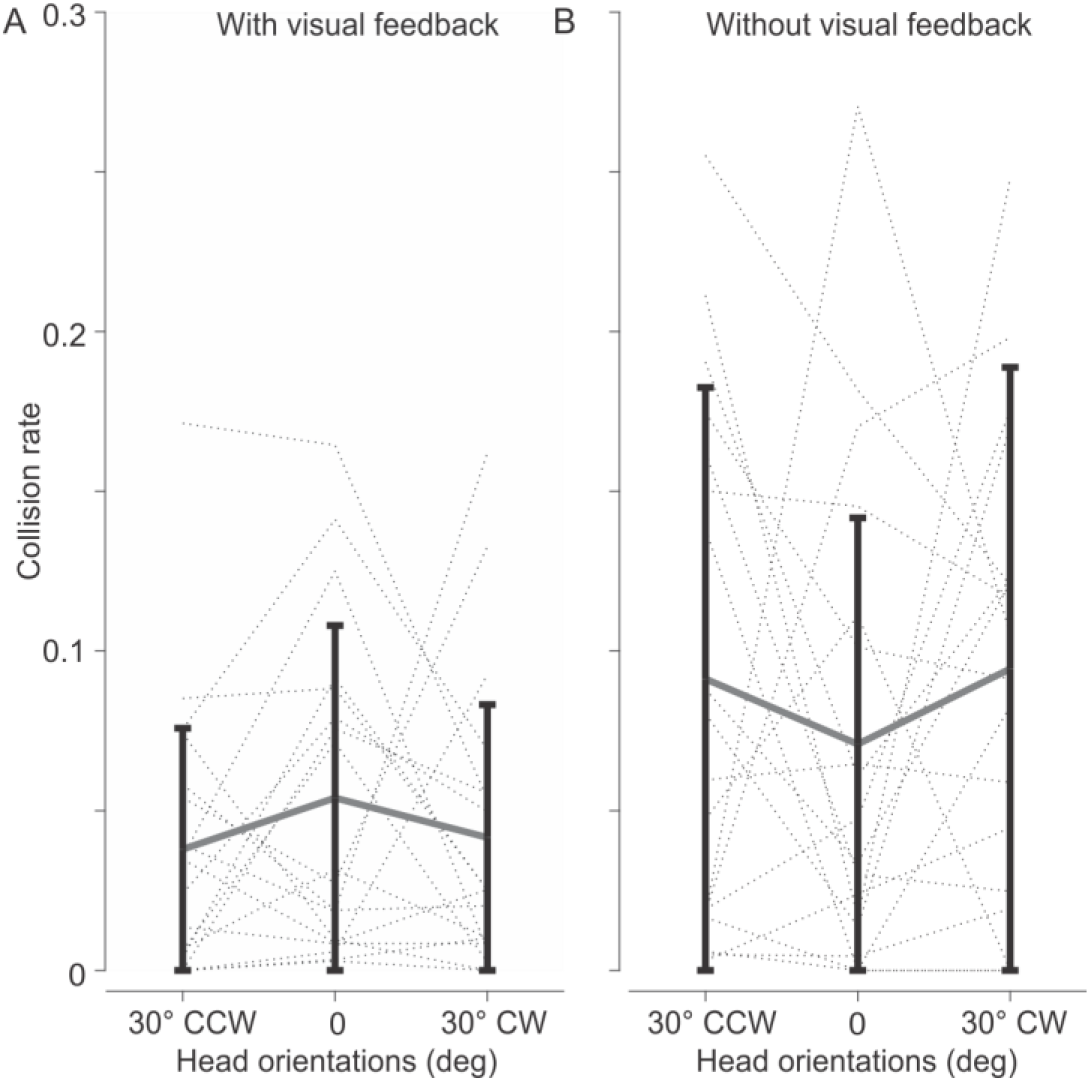
Effect of head orientation on collision rate. A) Visual feedback condition: participants were able to successfully perform the reaching task with a low collision rate across head orientations. B) No visual feedback condition: removing the visual feedback increased the overall collision rate irrespective of the head orientation. Error bars are standard deviations across participants.

### Participants adapted their obstacle avoidance behavior for head roll conditions

To further explore the effect of head roll on movement strategies, we determined the following parameters: movement speed, preferred direction of passing the obstacle, and expansion biases.

#### Movement speed

As mentioned before, reducing movement speed could be a compensation strategy to counteract the increased movement variability caused by rolling the head. However, we did not find any changes in movement speed for any of the experimental conditions (all p > 0.1).

#### Preferred direction of passing the obstacle

Figure 6 depicts the percentage of rightward movements for different head orientations, obstacle positions, and visual feedback conditions. Varying the head roll changed the preference in passing the obstacle from a certain side (left vs right). Rolling the head CCW led to a tendency to pass the obstacle from the right side, while rolling the head CW changed the tendency to pass the obstacle from the left side. Unsurprisingly, shifting the obstacle to the right or left of the central position changed the preferred direction of the movement. For example, when the centrally placed obstacle was shifted to the right, participants preferred to pass it from its left side and vice versa. Lastly, visual feedback of the movement did not seem to influence the passing side. The 3 (head orientation) × 5 (obstacle position) × 2 (visual feedback) repeated measures ANOVA on the percentage of rightward movements revealed a main effect of head orientation (F(2,34) = 12.564, p = 8.215e-4, η^2^ = 0.43), a main effect of obstacle position (F(4,68) = 290.279, p = 4.508e-21, η^2^=0.95), and an interaction between head orientation and obstacle position (F(8,136) = 405.711, p = 1.873e-4, η^2^ = 0.29). As can been seen in Figure 6 and revealed from the statistical analysis, there is no difference between the two obstacle configurations on the left (most leftward and leftward) or on the right of the central obstacle (most rightward and rightward). As there was no effect of visual feedback (p = 0.963) and no interaction between visual feedback and any other conditions (all p’s > 0.2), we collapsed the percentage of rightward movements across the visual feedback conditions as well as across the two leftward and the two rightward obstacle configurations (Figure 6C). The repeated measure ANOVA for the collapsed data for the central obstacle revealed a main effect of head orientation (F(2,68) = 15.91, p = 1.341e-5, η^2^ = 0.48). Post-hoc t-tests showed a significant difference between the three different head orientations (0° and 30° CW: t(17) = 3.076, p = 0.021, Cohen’s d = 0.73; 0° and 30° CCW: t(17) = 2.589, p = 0.019, Cohen’s d = 0.61; 30° CW and 30° CCW: t(17) = 5.703, p = 7.874e-6, Cohen’s d = 1.344). These results demonstrate that participants opted for more rightward and leftward passing movement when rolling the head CCW and CW compared to the straight head, respectively.

**Figure 6.**
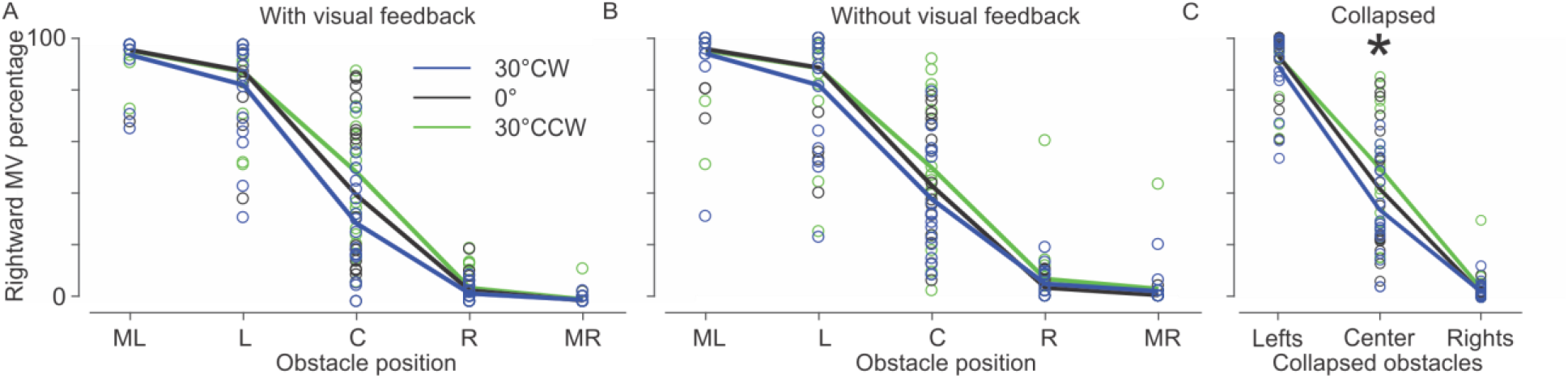
Percentage of rightward movements. Head roll caused changes in the preferred direction of movement A) with and B) without visual feedback. In both conditions rolling the head CCW increased the tendency in passing the obstacle from the right side. Obstacle positions: ML: most leftward, L: leftward, C: central, R: rightward, MR: most rightward. Open circles represent data from single participants. C) Since there was no difference between the leftward obstacle shifts (ML and L) and between the rightward obstacle shifts (R and ML), we collapsed the leftward and rightward shifts. The star indicates statistical significance with p < 0.05.

#### Safety margins

In addition to changing the preferred direction of passing the obstacle, increasing the safety margins by increasing the trajectory curvature could also compensate for the increased variability caused by head roll (as shown in Figure 2B). First, we had to select the trajectories for which we had enough data: we chose conditions with central obstacles as well as with non-central obstacles in which more than 10% of the overall movements passed from the same side (Figure 6). To remind the reader, we also expected to observe rotational biases caused by biases in coordinate transformations (Figure 2C). To demonstrate the effect of rotational and expansion biases on trajectories, we plotted the pooled trajectories for the central obstacle for both rightward and leftward movements and without visual feedback (Figure 7A). As can be seen, there is a symmetry between the shifts in trajectories for CW and CCW head rotations (zoomed left and right panels in Figure 7A) which is comparable to Figure 2D. The results show that rolling the head created both rotational (blue and green trajectories are shifted in opposite side of the black) and expansion (the trajectory shifts are not symmetrical; i.e. green trajectory being close to black one for rightward movements) biases.

**Figure 7.**
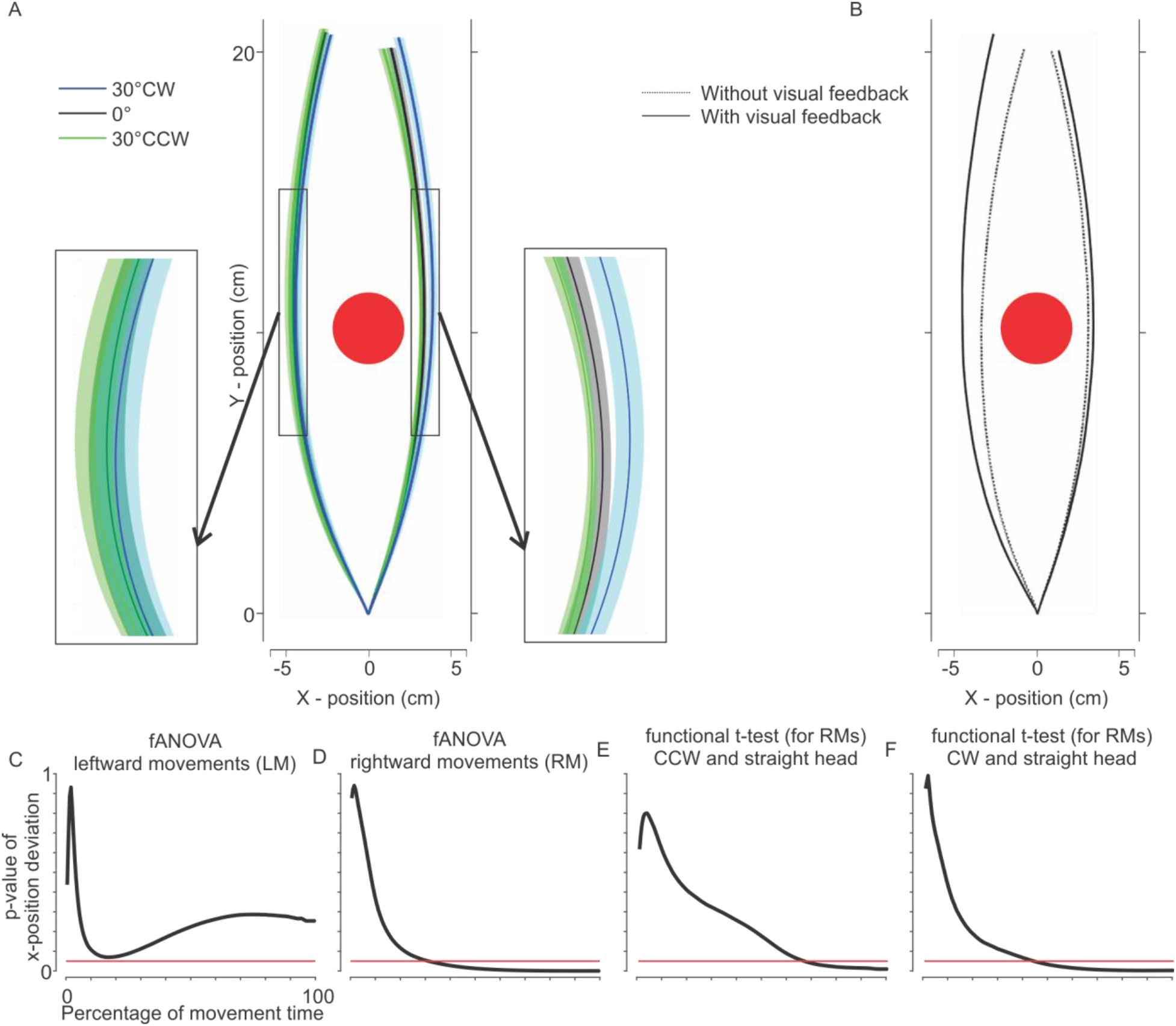
Participants showed both rotational and expansion biases caused by varying head orientation. A) Movement trajectories for the central obstacle with different head orientations in the absence of visual feedback. Trajectories are averaged across participants. The shaded areas indicate the standard errors of the mean. The obstacle is depicted as the red circle. Zoomed versions of the trajectories are provided, in two small panels, for better illustration of the effects. B) Comparison of the leftward and rightward movements in the presence (dotted lines) and absence (solid lines) of visual feedback. Removing visual feedback caused significant shifts of the trajectory for leftward movements but not for rightward movements. C and D: Repeated measure fANOVA for leftward and rightward movements. The red line indicates the p< 0.05. C) Different head rolls led to similar trajectories for passing the central obstacle on the left side. D) Different head rolls led to deviations of the trajectories for passing the central obstacle from the right side. E and F: Repeated measure functional comparison (t-test) of trajectories for rightward movements. E) Trajectories for straight and CCW head rolls overlapped except around the end of the movement. F) Trajectories for straight and CW head rolls deviated around 40% of movement time and later (before passing the obstacle).

The effect of head roll was more noticeable for rightward (Figure 7A; zoomed panel on the right) compared to leftward movements (Figure 7A; zoomed panel on the left). However, one needs to consider that the shift caused by removing the visual feedback for leftward movements while the head was straight was already stronger than for the rightward movements (Figure 7B; see the difference between the black solid line - without visual feedback and the black dotted line - with visual feedback. This observation is in line with previous studies (Chapman & Goodale, 2008; De Haan et al., 2014; Menger, Dijkerman, et al., 2013; Menger et al., 2012; Ross et al., 2018; Ross et al., 2015), and as Menger et al. (2013) illustrated, this is mainly due to the degree of obstructiveness of the obstacle. This indicates that people adapt their compensatory behavior if it is necessary. In other words, if the safety margins for the leftward movements were sufficiently large, there is no further need to increase the margins in the presence of higher uncertainty.

To assess if the deviation in trajectories for different head rolls are statistically significant, we deploy functional analysis of variability (fANOVA). A fANOVA provides F-statistics of where the continuous trajectories deviated from each other due to the experimental condition. We first ran the repeated fANOVA on time-normalized trajectories for different head roll conditions separately for each movement direction (Figure 7C-D). As can be seen the effect of the head roll on leftward movements (Figure 7C) never reaches significance while the head roll on rightward movements (Figure 7D) caused a deviation between trajectories after 30% of time passed. To further examine the effect of CW and CCW head roll on rightward movements, we ran function t-tests. The trajectory for CCW head orientation only deviated from the straight head trajectory toward the end of the reaching (long after passing the obstacle) (Figure 7E), while the CW head rotation caused an early deviation of the trajectory (before passing the obstacle) compared to the straight head trajectory (Figure 7F). This confirms the hypothesis illustrated in Figure 2D showing that both expansion and rotational biases are combined, with expansion biases being more noticeable around the time of passing the obstacle.

To quantify the effect of head orientation on safety margins, we first separated the rotational biases (due to misestimation of the head angle) from the expansion biases (due to uncertainty in head angle estimation). For details of this calculation please see “Materials and Methods” section. We performed the calculations for each individual participant and each obstacle, separately for each visual condition. Figure 8 illustrates expansion biases for the different conditions. Positive expansion biases indicate an increase in curvature (safety margins) and negative biases a decrease in curvature. This expansion or shrinkage is calculated compared to straight head conditions (see “Material and Method”). For the central obstacle, participants increased their safety margins only when they were passing the obstacle from the right side without visual feedback of their hand (t(13)=3, p = 0.01, Cohen’s d = 0.802) while they produced almost the same trajectories in all the other conditions (without visual feedback and leftward movement: t(13)=0.648, p = 0.528, Cohen’s d = 0.173; with visual feedback and rightward movement: t(13)=0.557, p = 0.587, Cohen’s d = 0.149; with visual feedback and leftward movement: t(13)=0.555, p = 0.589, Cohen’s d = 0.148). For obstacles shifted to the right (rightward and most rightward), we only considered leftward movements. While participants increased their curvature for the most rightward obstacle despite the presence or absence of visual feedback (with visual feedback: t(17)=3.675, p = 0.002, Cohen’s d = 0.866; without visual feedback: t(17)=2.291, p = 0.035, Cohen’s d = 0.540), they only did so for the rightward obstacle in the presence of visual feedback (t(17)=3.198, p = 0.005, Cohen’s d = 0.754) but not in the absence of visual feedback (t(16)=1.647, p = 0.119, Cohen’s d = 0.399). Similarly, for the obstacles shifted to the left we only considered rightward movements. We observed that both in the presence and absence of visual feedback participants significantly increased their curvature (most leftward and with visual feedback: t(17)=4.024, p = 8.805e-4, Cohen’s d = 0.948; most leftward and without visual feedback: t(17)=4.457, p = 3.462e-4, Cohen’s d = 1.051; leftward and with visual feedback: t(17)=2.982, p = 0.008, Cohen’s d = 0.703; leftward and without visual feedback: t(17)=2.468, p = 0.024, Cohen’s d = 0.582).

**Figure 8.**
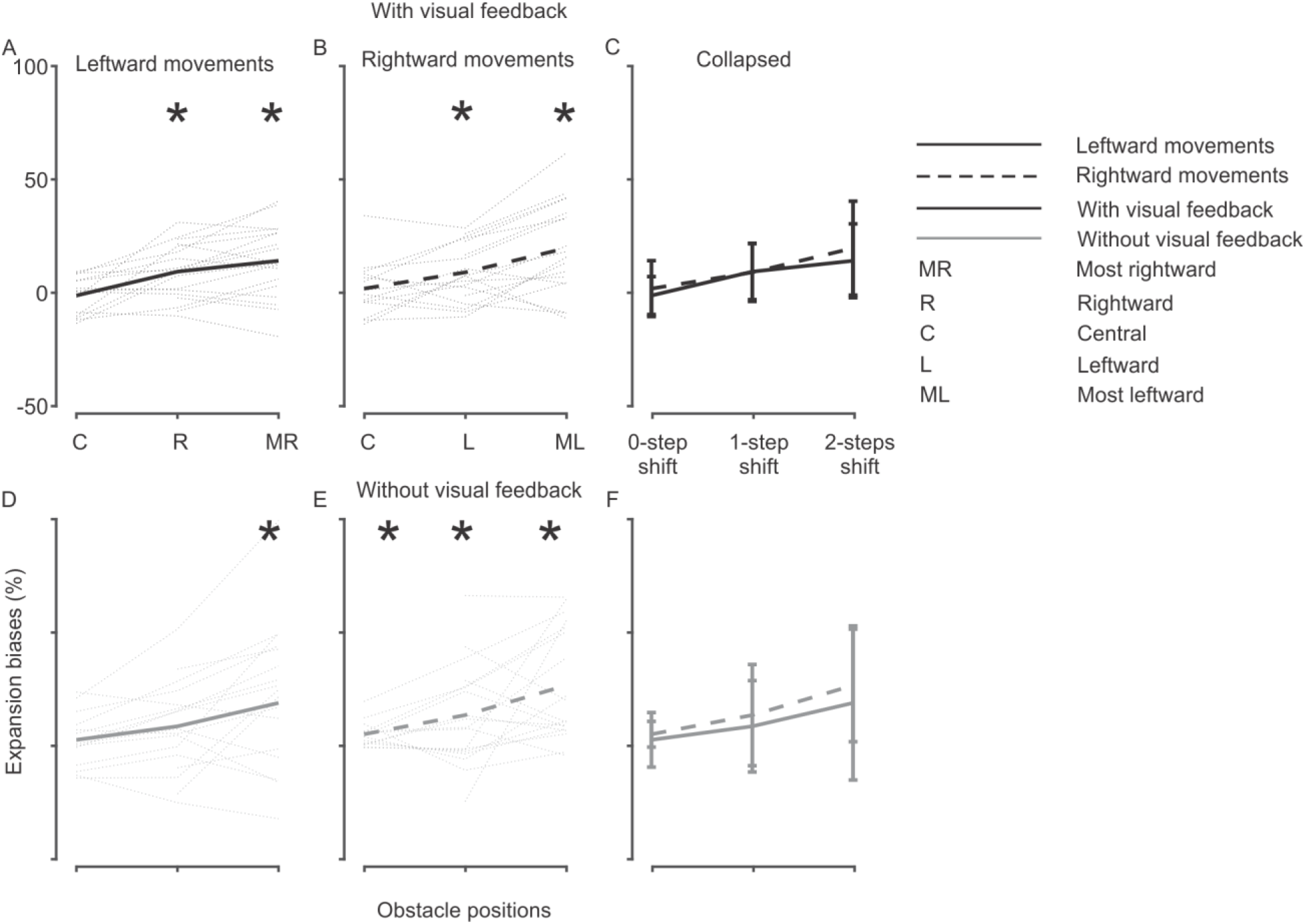
Rolling the head caused increasing safety margins dependent on the original curvature. A-C) Expansion biases for different obstacle positions and different movement directions in the presence of visual feedback. Dotted lines represent individual participants. A) Leftward movements: In order to assure that we have enough data points, we only considered leftward movements for the obstacles shifted to the right (R and MR). B) Rightward movements: similarly, we only considered rightward movements for the obstacles shifted to left (L and ML). C) Collapsed: to visually compare leftward and rightward movements we collapsed A and B. We renamed the obstacle positions for consistency: 0-step shift for the central obstacle position, 1-step shift for the rightward or leftward obstacle positions, and 2-steps shift for the most rightward or most leftward obstacle positions. D-F) Expansion biases for different obstacle positions and different movement directions in the absence of visual feedback. Conditions are identical to upper panels. Error bars represents standard error of the mean. Asterisks represent p < 0.05.

From the above analysis, we conclude that overall participants increased their safety margins for rolled head conditions compared to the straight head condition. However, this increase depends on the original curvature in the straight head condition. That is if the curvature for the straight head condition already provides sufficient safety margin, no further increase in safety margin is required for rolled head condition. In our task, the central obstacle is the most intruding obstacle and initially demands higher curvature compared to the shifted obstacle positions. Hereof, we observed no increase in curvature for the rolled head conditions in the presence of visual feedback. Analogously, since leftward movements were more curved in the absence of visual feedback, participants didn’t increase movement curvature for central and rightward obstacles (the two that were more intruding compared to most rightward obstacle).

## Discussion

The goal of the current study was to assess whether and how humans account for the added movement uncertainty induced by stochastic coordinate transformations in goal-directed movements. To this aim, we asked human participants to reach to visual targets while avoiding obstacles. In addition, we varied head orientations (straight and 30° CW/CCW) and visual feedback of the hand (with/without visual feedback). We hypothesized that if humans are compensating for the increased uncertainty caused by stochastic coordinate transformations, varying head orientation should not affect their performance (i.e. same collision rate for all head orientations). If that was true, we hypothesized to observe compensatory effects in the trajectories, such as increased safety margins (increased curvature), for the rolled compared to the straight head conditions. As expected, rolling the head increased movement variability. To accommodate this increased variability, participants adapted their movement behavior by varying their preferred movement direction (compared to the straight head condition) and increasing their safety margins from the obstacle (based on collision likelihood). Consequently, the collision rate remained the same for all head orientations. Thus, the human brain seems to consider the increased movement variability resulting from stochastic coordinate transformations when performing goal-directed movements.

The main assumption of the current study is that the stochasticity of coordinate transformations propagates to the final motor output. This assumption is based on numerous studies demonstrating that uncertainty in coordinate transformations causes higher variability in movement execution (Abedi Khoozani & Blohm, 2018; Biguer, Prablanc, & Jeannerod, 1984; Bock, 1986, 1993; Burns & Blohm, 2010; Henriques el al., 1998; Henriques & Crawford, 2000; Lewald & Ehrenstein, 2002; McGuire & Sabes, 2009; Schlicht & Schrater, 2007; Schütz, Henriques, & Fiehler, 2013; Sober & Sabes, 2003, 2005; Vaziri, Diedrichsen, & Shadmehr, 2006). For instance, when reaching to visual targets of different eccentricities with respect to gaze fixation, reaching movements overshoot the target in the absence of visual feedback (Bock, 1986, 1993; Henriques et al., 1998; Henriques & Crawford, 2000; Lewald & Ehrenstein, 2002; Vaziri et al., 2006). It has been suggested that this overshoot likely arises from noise in transforming the visual estimate of the target into the proprioceptive estimate of the hand (Dessing et al., 2012). Furthermore, McGuire and Sabes (2009) showed that the gaze-dependent errors vary based on the target’s modality (visual or proprioceptive or both) as well as the available information of the initial hand position (with or without visual feedback). They found that gaze-dependent reaching errors are only observable for visual targets and are abolished for proprioceptive targets, suggesting that the transformation of a visual target into the coordinate frame of the arm systematically affects reaching movements. Based on the evidence that accurate coordinate transformations rely on the estimation of body geometry (Blohm & Crawford, 2007) and to elaborate on the effect of stochastic coordinate transformations on reaching movements, previous studies varied the reliability of the head angle estimations via rolling the head and/or loading the neck (Abedi Khoozani & Blohm, 2018; Burns & Blohm, 2010). Both factors biased reaching movements and increased the movement variability compared to a control condition (e.g. straight head and no neck load). Given these observations, we argue that there is a clear propagation of uncertainty caused by stochastic coordinate transformations to the performed reaching movements.

If stochastic coordinate transformations cause higher movement variability, does the brain account for such noise when planning and executing goal-directed movements? In the following, we argue that it is rather unlikely that the brain dismisses such nuisances.

We observed two main strategies to compensate for the increased movement variability caused by stochastic coordinate transformations: (1) changes in the preferred movement direction when passing the obstacle, and (2) increased safety margins.

With regards to strategy (1), we believe that it is caused by signal-dependent noise. Since for the rightward/leftward obstacles one direction is distinctly dominant (e.g. rightward direction for the obstacle shifted to the most leftward positions), we only focus on the central obstacle in which the likelihood of passing the obstacle from the right- or left side was (almost) at chance level. To elaborate more on why changing the preferred direction will facilitate the effect of coordinate transformations we exemplarily consider the 30° CW head orientation. In this configuration, participants preferred to pass the central obstacle from the left side, while they passed it from the right side when the head was straight (Figure 6). We believe that participant might have changed their preferred movement direction to avoid extra rotation-translation of their eyes. It has been shown that humans move their gaze to specific task related landmarks (e.g. possible contact point with obstacle) during reaching movements to gain spatial information for movement control (Johansson et al., 2001). Using this analogy, for 30° CW head orientation a rightward eye rotation is required in order to have a more accurate view of the right side of the screen (with regard to the body and screen midpoint) while such rotation is not required for left side of the screen. Meanwhile, the accurate coordinate transformation relies on the estimation of body geometry (Blohm & Crawford, 2007), here both head and eye angles. Additionally, it has been established that the noise associated with this estimation is signal-dependent (Abedi Khoozani & Blohm, 2018; Burns & Blohm, 2010; Schlicht & Schrater, 2007); that is, the larger the amplitude of the signal, the noisier the estimation. Therefore, the extra rotation-translation of the eyes, required for the right side of the screen, may result in noisier eye-in-head orientation estimations and, consequently, noisier coordinate transformations. Accordingly, to decrease the uncertainty associated with the coordinate transformation it may be beneficial to pass the obstacle on the left side, which is also in accordance with our data (see Figure 6C). Hence, we argue that a likely explanation for the change of passing direction as a function of head roll is that participants systematically adapted their preferred movement direction to decrease the likelihood of hitting the obstacle that arises from the uncertainty accompanied by the required coordinate transformations.

With regards to strategy (2), we observed increased safety margins for non-central obstacles in both the presence and absence of visual feedback. While the appearance of expansion biases in the absence of visual feedback is expected, the persistence of expansion biases in the presence of visual feedback is somewhat interesting. This is mainly due to the observation that providing visual feedback of the hand will remove the biases caused by gaze-shifts and more generally by coordinate transformations (Brown, Marlin, & Morrow, 2015; Dessing et al., 2012; Saunders, 2004; Saunders & Knill, 2003). However, our results suggest that the signal-dependent noise still persists in the system. In other words, while the extra source of information (i.e. visual information of the hand position) decreases the amount of uncertainty, it is not fully abolished. Similarly, Ross et al. (2015) observed that varying the fixation while visual feedback of the hand was available caused participants to veer away from the fixated obstacle as opposed to free viewing or central fixation. The authors speculated that the observed pattern can be explained by a misestimation of the target position on the retina, however, we argue that the observed veering away from the fixated obstacle might be better explained by stochastic coordinate transformations: Given that varying gaze position will result in higher uncertainty in eye-in-head orientation estimation and consequently in noisier movements, it is logical to increase the safety margin to decrease the likelihood of obstacle collision.

Furthermore, we observed that the increase in the curvature for the rolled head conditions depends on the initial curvature during straight head trials. That is, for the obstacles shifted to the right or left participants increased their safety margins for rolled head conditions in both presence and absence of visual feedback. We believe that this can be explained by considering the biomechanical constraints of the performing limb. Numerous studies demonstrated that the central nervous system incorporates biomechanics (kinematics and dynamics) of the moving limb in planning and executing reaching movements to visual targets (Cos, Duque, & Cisek, 2014; Nashed, Crevecoeur, & Scott, 2012; Sabes & Jordan, 1997; Sabes, Jordan, & Wolpert, 1998). More specifically, it has been shown that humans are capable of estimating the possible biomechanical costs of their trajectories very early in movement planning (200ms; for details refer to Cos et al., 2014) and select the trajectory with the lowest biomechanical cost which satisfies the task requirements (Cos et al., 2014). In addition to biomechanical costs, it has been shown that in obstacle avoidance tasks, humans align their trajectories to achieve lower sensitivity to position uncertainties or possible perturbations of the performing limb (Nashed, Crevecoeur, & Scott, 2012; Sabes & Jordan, 1997; P. N. Sabes, Jordan, & Wolpert, 1998; Voudouris et al., 2012). For instance, Voudouris et al. (2012) showed that when humans are passing an obstacle, they accurately consider the possibility of hitting the obstacle with their whole moving arm and change their upper-limb configuration to accommodate the task. In our data, this is specifically observable for the rightward and leftward trajectories in the absence of visual feedback (Figure 7B). That is, as all participants performed their movements with their right hand, passing obstacles from the left side requires higher curvature to avoid any possible collision with the performing arm (as opposed to only considering the fingertips or hand). This is in accordance with what has been reported in many previous studies (Chapman & Goodale, 2008; De Haan et al., 2014; Menger et al., 2013; Menger et al., 2012; Ross et al., 2018; Ross et al., 2015; Voudouris et al., 2012). Hence, together with previous findings, our results suggest that humans are trading of between minimizing the likelihood of obstacle collision (by increasing the curvature) and minimizing the biomechanical costs (by decreasing the curvature).

As mentioned before, we believe that increasing the safety margin is a strategy to account for the signal dependent noise caused by stochastic coordinate transformations. Therefore, it is crucial to investigate how signal dependent noise affects the calculation of collision likelihood and motor planning. This has been studied previously (Hamilton & Wolpert, 2002; Harris & Wolpert, 1998). For instance, Harris and Wolpert (1998) proposed a theoretical framework, called task optimization in the presence of signal-dependent noise (TOPS), and showed that including signal-dependent noise provides a general framework which can explain both saccadic and point-to-point arm movements. In a later study, Hamilton and Wolpert (2002) extended this framework and showed that TOPS can also predict the trajectories generated in an obstacle avoidance task. Based on these observations, they proposed that controlling the statistics of the movement (such as minimizing the endpoint error) while accounting for signal-dependent noise might offer a unifying principle for goal-directed movements. Therefore, it is also crucial to have a better understanding of the nature of such signal-dependent noise corrupting the movements. While previous studies (Hamilton & Wolpert, 2002; Harris & Wolpert, 1998; Van Beers, Baraduc, & Wolpert, 2002) mainly associated the signal-dependent noise with the amplitude of the motor command, we argue that the processing, e.g. coordinate transformations, required for generating the motor command can also cause movement variability and therefore, it is important to account for such noise in the motor circuitry.

While the role of coordinate transformations in the motor planning stage is shown through many studies (Abedi Khoozani & Blohm, 2018; Burns & Blohm, 2010; McGuire & Sabes, 2009; Sober & Sabes, 2003, 2005), its impact in the optimal motor control field is less understood. This study provided the first evidence that the brain accommodates for the added movement variability due to uncertainty in coordinate transformations. However, it is not clear at what stage during motor planning and execution such accommodations might occur. Based on the optimal control theory, the motor system selects the appropriate control law to calculate the motor command based on the desired task goal (e.g. grab a pen) and the current system states (i.e. limb position). Since both motor commands and sensory signals of motor performance are corrupted with noise, the optimal state estimator uses both sensory signals (feedback circuitry) and an efference copy of the motor commands (feedforward circuitry) to estimate the current state of the limb (Scott & Norman, 2003; Scott, 2004; Todorov, 2004; Todorov & Jordan, 2002). Within this framework, however, it is unknown in which coordinates each of these processes (feedback and feedforward) can be carried out. It has been shown that it is beneficial to plan movements in multiple coordinate frames (McGuire & Sabes, 2009). In addition, the feedback of the movement can be presented in different coordinate frames, e.g. the visual feedback of the hand in retinal coordinates and the proprioceptive feedback of the hand in body coordinates. Thus, it is not trivial which coordinate system should be used for implementing optimal feedback control. For instance, all signals could be transformed and then combined in one coordinate frame (e.g. visual, proprioceptive, or an intermediate frame). Alternatively, all signals could be transformed into the other signal’s coordinate frame (i.e. visual to proprioceptive and vice versa) and the error signal is generated in all coordinate frames in parallel (similar to generating movement plans in multiple coordinate systems). The exact role of coordinate transformations in motor control circuit requires further investigation.

On the other hand, the brain might switch from an optimal control strategy to a model-free control strategy (i.e. robust control) to account for the disturbances caused by head roll (Crevecoeur et al., 2019). This is based on the new findings that when participants encountered with unpredictable disturbances (i.e. sudden force field applied to the moving hand), they deployed robust control strategy (e.g. faster movement and more rigorous response to disturbances). However, when disturbances were predictable, participants deployed strategies closer to optimal feedback control. Whether the uncertainties induced due to head roll can cause a switch in control strategy is not known currently. Further modeling and experimental studies are required to investigate the role of coordinate transformations in the optimal motor control framework. Such studies have implications not only in the motor control field, but also in perception, decision making as well as applicable fields such as brain machine interfaces and robotics.

All in all, we believe that uncertainty in coordinate transformations stemming from signal-dependent noise propagates to motor behavior and that the brain accounts for such noise during motor planning, and possibly execution.

## Acknowledgements

We would like to thank Marie Mosebach for her help in data collection. This project was funded by NSERC (Canada), DFG IRTG 1901 “The brain in action” (Germany), and by scholarship to PAK by the German Academic Exchange Service (DAAD).

## ENDNOTE

At the request of the author(s), readers are herein alerted to the fact that additional materials related to this manuscript may be found at [Data: https://osf.io/tf8p5/ and analysis codes: https://github.com/Parisaabedi/Obstacle-Avoidance]. These materials are not a part of this manuscript and have not undergone peer review by the American Physiological Society (APS). APS and the journal editors take no responsibility for these materials, for the website address, or for any links to or from it.

## Supplementary Material

**Figure S1.**
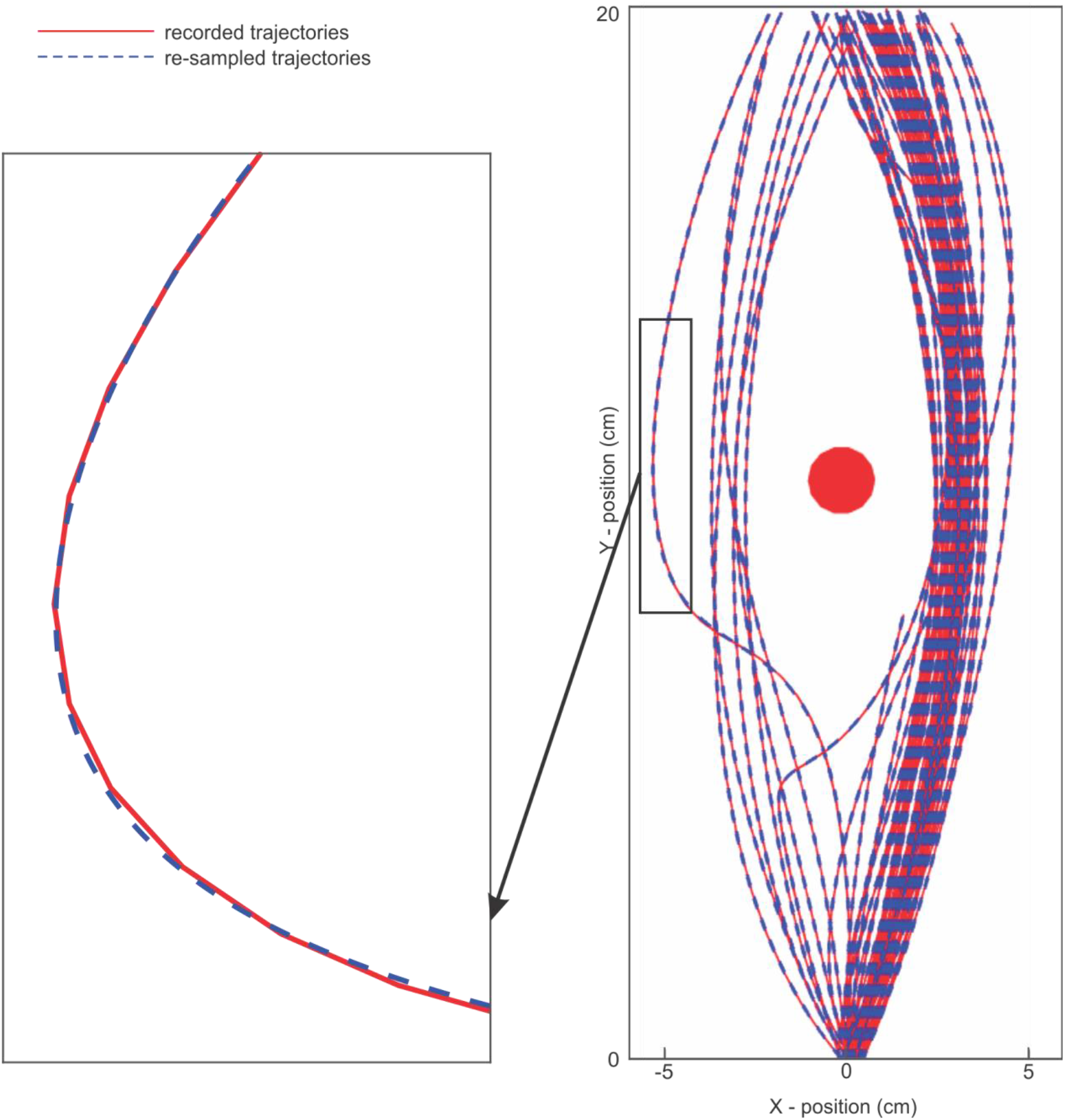
Comparison between re-sampled and recorded trajectories. Right panel shows several trajectories (resampled and recorded) for the central obstacle. The left panel shows a zoomed illustration of an example trajectory. As can be seen, the algorithm generated an acceptable resemblance of the recorded trajectories.

## References

Abedi Khoozani, P., & Blohm, G. (2018). Neck muscle spindle noise biases reaches in a multisensory integration task. Journal of Neurophysiology, 120(3), 893–909. https://doi.org/10.1152/jn.00643.2017

Alikhanian, H., Carvalho, S. R., & Blohm, G. (2015). Quantifying effects of stochasticity in reference frame transformations on posterior distributions. Frontiers in Computational Neuroscience, 9, 82. https://doi.org/10.3389/fncom.2015.00082

Battaglia, P. W., & Schrater, P. R. (2007). Humans Trade Off Viewing Time and Movement Duration to Improve Visuomotor Accuracy in a Fast Reaching Task. Journal of Neuroscience, 27(26), 6984–6994. https://doi.org/10.1523/JNEUROSCI.1309-07.2007

Biguer, B., Prablanc, C., & Jeannerod, M. (1984). The contribution of coordinated eye and head movements in hand pointing accuracy. Experimental Brain Research, 55, 462–469.

Blohm, G., & Crawford, J. D. (2007). Computations for geometrically accurate visually guided reaching in 3-D space. Journal of Vision, 7(5), 4. https://doi.org/10.1167/7.5.4

Bock, O. (1986). Contribution of retinal versus extraretinal signals towards visual localization in goal-directed movements. Experimental Brain Research, 64, 476–482.

Bock, O. (1993). Localization of objects in the peripheral visual field. Behavioural Brain Research, 56(1), 77–84. https://doi.org/10.1016/0166-4328(93)90023-J

Brenner, E., & Smeets, J. B. J. (2018). How Can You Best Measure Reaction Times? Journal of Motor Behavior, 25. https://doi.org/10.1080/00222895.2018.1518311

Brown, L. E., Marlin, M. C., & Morrow, S. (2015). On the contributions of vision and proprioception to the representation of hand-near targets. Journal of Neurophysiology, 113(2), 409–19. https://doi.org/10.1152/jn.00005.2014

Buneo, C. A., & Andersen, R. A. (2006). The posterior parietal cortex: sensorimotor interface for the planning and online control of visually guided movements. Neuropsychologia, 44(13), 2594–606. https://doi.org/10.1016/j.neuropsychologia.2005.10.011

Buneo, C. A., Jarvis, M. R., Batista, A. P., & Andersen, R. A. (2002). Direct visuomotor transfomrations for reaching. Nature, 416(3), 632–636.

Burns, J. K., & Blohm, G. (2010). Multi-sensory weights depend on contextual noise in reference frame transformations. Frontiers in Human Neuroscience, 4, 221. https://doi.org/10.3389/fnhum.2010.00221

Burns, J. K., Nashed, J. Y., & Blohm, G. (2011). Head roll influences perceived hand position. Journal of Vision, 11(9), 3. https://doi.org/10.1167/11.9.3

Chapman, C. S., & Goodale, M. A. (2008). Missing in action: The effect of obstacle position and size on avoidance while reaching. Experimental Brain Research, 191(1), 83–97. https://doi.org/10.1007/s00221-008-1499-1

Chua, R., & Elliott, D. (1993). Visual regulation of manual aiming. Human Movement Science, 12(4), 365–401. https://doi.org/10.1016/0167-9457(93)90026-L

Crevecoeur, F., Scott, S.H., Cluff, T., (2019) Robust control in human reaching movements: a model-free strategy to compensate for unpredictable disturbances. Journal of Neuroscience. PMID: 31488611 DOI: 10.1523/JNEUROSCI.0770-19.2019

Cohen, R. G., Biddle, J. C., & Rosenbaum, D. A. (2010). Manual obstacle avoidance takes into account visual uncertainty, motor noise, and biomechanical costs. Experimental Brain Research, 201(3), 587–592. https://doi.org/10.1007/s00221-009-2042-8

Cohen, Y. E., & Andersen, R. A. (2002). A common reference frame for movement plans in the posterior parietal cortex. Nature Reviews Neuroscience, 3(7), 553–562. https://doi.org/10.1038/nrn873

Cos, I., Duque, J., & Cisek, P. (2014). Rapid prediction of biomechanical costs during action decisions. Journal of Neurophysiology, 112(6), 1256–1266. https://doi.org/10.1152/jn.00147

De Haan, A. M., Van der Stigchel, S., Nijnens, C. M., & Dijkerman, H. C. (2014). The influence of object identity on obstacle avoidance reaching behaviour. Acta Psychologica, 150, 94–99. https://doi.org/10.1016/j.actpsy.2014.04.007

Dessing, J. C., Byrne, P. A., Abadeh, A., & Crawford, J. D. (2012). Hand-related rather than goal-related source of gaze-dependent errors in memory-guided reaching. Journal of Vision, 12(11), 17–17. https://doi.org/10.1167/12.11.17

Engel, K. C., Flanders, M., & Soechting, J. F. (2002). Oculocentric frames of reference for limb movement. Archives Italiennes de Biologie. 140(3), 211–219.

Gallivan, J. P., & Chapman, C. S. (2014). Three-dimensional reach trajectories as a probe of real-time decision-making between multiple competing targets. Frontiers in Neuroscience, 8(8 JUL), 1–19. https://doi.org/10.3389/fnins.2014.00215

Hamilton, A. F. D. C., & Wolpert, D. M. (2002). Controlling the statistics of action: obstacle avoidance. Journal of Neurophysiology, 87(5), 2434–2440. https://doi.org/10.1152/jn.00875.2001.

Harris, C. M., & Wolpert, D. M. (1998). Signal-dependent noise determines motor planning. Nature, 394(6695), 780–784. https://doi.org/10.1038/nature03031.1.

Heath, M. (2005). Role of Limb and Target Vision in the Online Control of Memory-Guided Reaches. Motor Control, 9, 281–309. https://doi.org/10.1123/mcj.9.3.281

Heath, M., Westwood, D. A., & Binsted, G. (2004). The Control of Memory-Guided Reaching Movements in Peripersonal Space. Motor Control, 8, 76–106. https://doi.org/10.1123/mcj.8.1.76

Henriques, D. Y., Klier, E. M., Smith, M. A., Lowy, D., & Crawford, J. D. (1998). Gaze-centered remapping of remembered visual space in an open-loop pointing task. Journal of Neuroscience, 18(4), 1583–94. Retrieved from http://www.ncbi.nlm.nih.gov/pubmed/9454863

Henriques, D. Y. P., & Crawford, J. D. (2000). Direction-dependent distortions of retinocentric space in the visuomotor transformation for pointing. Experimental Brain Research, 132(2), 179–194. https://doi.org/10.1007/s002210000340

Howard, I. S., Ingram, J. N., & Wolpert, D. M. (2009). A modular planar robotic manipulandum with end-point torque control. Journal of Neuroscience Methods, 181(2), 199–211. https://doi.org/10.1016/j.jneumeth.2009.05.005

Jacobson, M., & Matthews, P. (1998). Generating uniformly distributed random Latin squares. Journal of Combinatorial Designs, 4(6), 405–437. https://doi.org/10.1002/(SICI)1520-6610(1996)4:6<405::AID-JCD3>3.0.CO;2-J

Khan, M. A., Elliott, D., Coull, J., Chua, R., & Lyons, J. (2002). Optimal control strategies under different feedback schedules: Kinematic evidence. Journal of Motor Behavior, 34(1), 45–57. https://doi.org/10.1080/00222890209601930

Knudsen, E. (2002). Computational Maps In The Brain. Annual Review of Neuroscience, 10, 41–65. https://doi.org/10.1146/annurev.neuro.10.1.41

Knudsen, E. I., du Lac, S., & Esterly, S. D. (2002). Computational Maps In The Brain. Annual Review of Neuroscience, 10, 41–65. https://doi.org/10.1146/annurev.neuro.10.1.41

Lacquaniti, & Caminiti. (2003). Visuo-motor transformations for arm reaching. European Journal of Neuroscience, 10(1), 195–203. https://doi.org/10.1046/j.1460-9568.1998.00040.x

Lewald, J., & Ehrenstein, W. H. (2002). Auditory-visual spatial integration: A new psychophysical approach using laser pointing to acoustic targets. The Journal of the Acoustical Society of America, 104(3), 1586–1597. https://doi.org/10.1121/1.424371

Loehr, J. D., & Palmer, C. (2007). Cognitive and biomechanical influences in pianists’ finger tapping. Experimental Brain Research, 178(4), 518–528. https://doi.org/10.1007/s00221-006-0760-8

McGuire, L. M. M., & Sabes, P. N. (2009). Sensory transformations and the use of multiple reference frames for reach planning. Nature Neuroscience, 12(8), 1056–1061. https://doi.org/10.1038/nn.2357

Menger, R., Dijkerman, H. C., & Van der Stigchel, S. (2013). The Effect of Similarity: Non-Spatial Features Modulate Obstacle Avoidance. PLoS ONE, 8(4). https://doi.org/10.1371/journal.pone.0059294

Menger, R., Van der Stigchel, S., & Dijkerman, H. C. (2012). How obstructing is an obstacle? The influence of starting posture on obstacle avoidance. Acta Psychologica, 141(1), 1–8. https://doi.org/10.1016/j.actpsy.2012.06.006

Nashed, J. Y., Crevecoeur, F., & Scott, S. H. (2012). Influence of the behavioral goal and environmental obstacles on rapid feedback responses. Journal of Neurophysiology, 108(4), 999–1009. https://doi.org/10.1152/jn.01089.2011

Ramsay, J., & Silverman, B. W. (2005). Functional Data Analysis. New York, NY: Springer. https://doi.org/10.1007/b98888

Ross, A. I., Schenk, T., Billino, J., Macleod, M. J., & Hesse, C. (2018). Avoiding unseen obstacles: Subcortical vision is not sufficient to maintain normal obstacle avoidance behaviour during reaching. Cortex, 98, 177–193. https://doi.org/10.1016/j.cortex.2016.09.010

Ross, A. I., Schenk, T., & Hesse, C. (2015). The effect of gaze position on reaching movements in an obstacle avoidance task. PLoS ONE, 10(12). https://doi.org/10.1371/journal.pone.0144193

Sabes, philip N. (1997). Obstacle Avoidance and a Perturbation Sensitivity Model for Motor Planning Philip. The Journal of Neuroscience, 17(18), 7119–7128.

Sabes, P. N., Jordan, M. I., & Wolpert, D. M. (1998). The Role of Inertial Sensitivity in Motor Planning. The Journal of Neuroscience, 18(15), 5948–5957. https://doi.org/10.1523/jneurosci.18-15-05948.1998

Saunders, J. A. (2004). Visual Feedback Control of Hand Movements. Journal of Neuroscience, 24(13), 3223–3234. https://doi.org/10.1523/jneurosci.4319-03.2004

Saunders, J. A., & Knill, D. C. (2003). Humans use continuous visual feedback from the hand to control fast reaching movements. Experimental Brain Research, 152(3), 341–352. https://doi.org/10.1007/s00221-003-1525-2

Schlicht, E. J., & Schrater, P. R. (2007). Impact of coordinate transformation uncertainty on human sensorimotor control. Journal of Neurophysiology, 97(6), 4203–4214. https://doi.org/10.1152/jn.00160.2007

Scott, S. H. (2004). Optimal feedback control and the neural basis of volitional motor control. Nature Reviews Neuroscience, 5(7), 532–545. https://doi.org/10.1038/nrn1427

Scott, S. H., & Norman, K. E. (2003). Computational approaches to motor control and their potential role for interpreting motor dysfunction. Current Opinion in Neurology. https://doi.org/10.1097/01.wco.0000102631.16692.71

Sober, S. J., & Körding, K. P. (2012). What silly postures tell us about the brain. Frontiers in Neuroscience, 6, 5–6. https://doi.org/10.3389/fnins.2012.00154

Sober, S. J., & Sabes, P. N. (2003). Multisensory integration during motor planning. The Journal of Neuroscience, 23(18), 6982–92. Retrieved from http://www.pubmedcentral.nih.gov/articlerender.fcgi?artid=2538489&tool=pmcentrez&rendertype=abstract

Sober, S. J., & Sabes, P. N. (2005). Flexible strategies for sensory integration during motor planning. Nature Neuroscience, 8(4), 490–7. https://doi.org/10.1038/nn1427

Soechting, J. F., & Flanders, M. (1992). Moving in three dimensional space: frames of reference, vectors, and coordinate systems. Annual Review of Neuroscience, 167–191.

Todorov, E. (2004). Optimality principles in sensorimotor control. Nature Neuroscience, 7(9), 907–915. https://doi.org/10.1038/nn1309

Todorov, E., & Jordan, M. I. (2002). Optimal feedback control as a theory of motor coordination. Nature Neuroscience, 5(11), 1226–1235. https://doi.org/10.1038/nn963

Van Beers, R. J., Baraduc, P., & Wolpert, D. M. (2002). Role of uncertainty in sensorimotor control. Philosophical Transactions of the Royal Society B: Biological Sciences, 357(1424), 1137–1145. https://doi.org/10.1098/rstb.2002.1101

Vaziri, S., Diedrichsen, J., & Shadmehr, R. (2006). Why Does the Brain Predict Sensory Consequences of Oculomotor Commands? Optimal Integration of the Predicted and the Actual Sensory Feedback. Journal of Neuroscience, 26(16), 4188–4197. https://doi.org/10.1523/jneurosci.4747-05.2006

Voudouris, D., Smeets, J. B. J., & Brenner, E. (2012). Do obstacles affect the selection of grasping points? Human Movement Science, 31(5), 1090–1102. https://doi.org/10.1016/j.humov.2012.01.005

Whitwell RL, Goodale MA. (2013) Grasping without vision: Time normalizing grip aperture profiles yields spurious grip scaling to target size. Neuropsychologia 51: 1878–1887.

